# Kinetic analysis reveals the rates and mechanisms of protein aggregation in a multicellular organism

**DOI:** 10.1101/2020.08.13.249862

**Authors:** Tessa Sinnige, Georg Meisl, Thomas C. T. Michaels, Michele Vendruscolo, Tuomas P.J. Knowles, Richard I. Morimoto

## Abstract

The accumulation of insoluble protein aggregates containing amyloid fibrils has been observed in many different human protein misfolding diseases^1,2^, and their pathological features have been recapitulated in diverse model systems^3^. *In vitro* kinetic studies have provided a quantitative understanding of how the fundamental molecular level processes of nucleation and growth lead to amyloid formation^4^. However, it is not yet clear to what extent these basic biophysical processes translate to amyloid formation *in vivo*, given the complexity of the cellular and organismal environment. Here we show that the aggregation of a fluorescently tagged polyglutamine (polyQ) protein into µm-sized inclusions in the muscle tissue of living *C. elegans* can be quantitatively described by a molecular model where stochastic nucleation occurs independently in each cell, followed by rapid aggregate growth. Global fitting of the image-based aggregation kinetics reveals a nucleation rate corresponding to 0.01 h^-1^ per cell at 1 mM intracellular protein concentration, and shows that the intrinsic stochasticity of nucleation accounts for a significant fraction of the observed animal-to-animal variation. Our results are consistent with observations for the aggregation of polyQ proteins *in vitro*^5^ and in cell culture^6^, and highlight how nucleation events control the overall progression of aggregation in the organism through the spatial confinement into individual cells. The key finding that the biophysical principles associated with protein aggregation in small volumes remain the governing factors, even in the complex environment of a living organism, will be critical for the interpretation of *in vivo* data from a wide range of protein aggregation diseases.

## Main

Protein aggregation is a pathological hallmark of a wide range of neurodegenerative and systemic diseases^1,2^. Chemical kinetics has been a powerful tool to elucidate the microscopic steps of the aggregation pathway for a variety of disease-associated proteins and peptides *in vitro*, most notably the amyloid-β peptide associated with Alzheimer’s disease (AD)^7^. This approach has been invaluable to the design of small molecules that reduce the generation of potentially toxic oligomeric species^8,9^, to understand the action of molecular chaperones in inhibiting protein aggregation^10,11^, and to rationalize the efficacy of antibodies in clinical trials for AD^12^. This framework has been extended to include stochastic and spatial effects that control kinetics in small volumes (fL-pL)^13–15^, with the premise of being applicable to protein aggregation in living cells.

However, cells and organisms have evolved intricate protein homeostasis pathways to ensure correct protein folding and to suppress misfolding and aggregation^16,17^, and it has not yet been established whether a chemical kinetics approach is sufficient to describe protein aggregation *in vivo*. The fundamental question is whether the complex nature of the cellular and organismal environment induces a major remodelling of the aggregation network as studied *in vitro*, or whether the same biophysical principles remain dominant. The nematode *C. elegans* provides the high level of control and the tools necessary to perform a quantitative kinetic analysis and determine the mechanisms governing protein aggregation in a living animal. *C. elegans* has a well-defined anatomy, and the animals within a population are genetically identical. Perhaps the most beneficial feature of this animal model system is its optical transparency, allowing the aggregation of a fluorescently labelled protein to be directly visualised. Specifically, we take advantage of the *C. elegans* muscle tissue that corresponds to 95 physiologically identical cells, and propose that these cells can be quantitatively modelled as individual “test tubes” in which the deposition of expanded polyQ takes place by a mechanism of nucleated aggregation.

### Establishing a framework for *in vivo* protein aggregation kinetics

PolyQ-containing proteins have been studied extensively *in vitro* and *in vivo*, and expansions of polyQ were predicted^18^, and have been observed to form cross-β fibrils *in vitro*^19,20^ and to accumulate into insoluble inclusions in polyQ diseases *in vivo*^19,21,22^. When expressed in *C. elegans* muscle cells, a Q40 protein C-terminally tagged with yellow fluorescent protein (Q40-YFP) forms intracellular inclusions distributed throughout the tissue that have a relatively uniform size and shape (Fig. 1a, Extended Data Fig. 1)^23^. Visualised by transmission electron microscopy (TEM), the polyQ inclusions contain fibrillar material with dimensions on the order of 10 nm (Fig. 1b, Extended Data Fig. 2). The appearance of immobile amyloid-like inclusions is consistent with fluorescence lifetime imaging (FLIM)^24^ and fluorescence recovery after photobleaching (FRAP)^23^ experiments carried out previously. Protein aggregation kinetics can thus be monitored in living animals using the visualisation of intracellular Q40-YFP inclusions as a direct measure of the total aggregate amount. The appearance of inclusions over time is reminiscent of the kinetics of amyloid formation observed *in vitro*, displaying a rapid increase from zero towards a plateau (Fig. 1c).

**Fig. 1:**
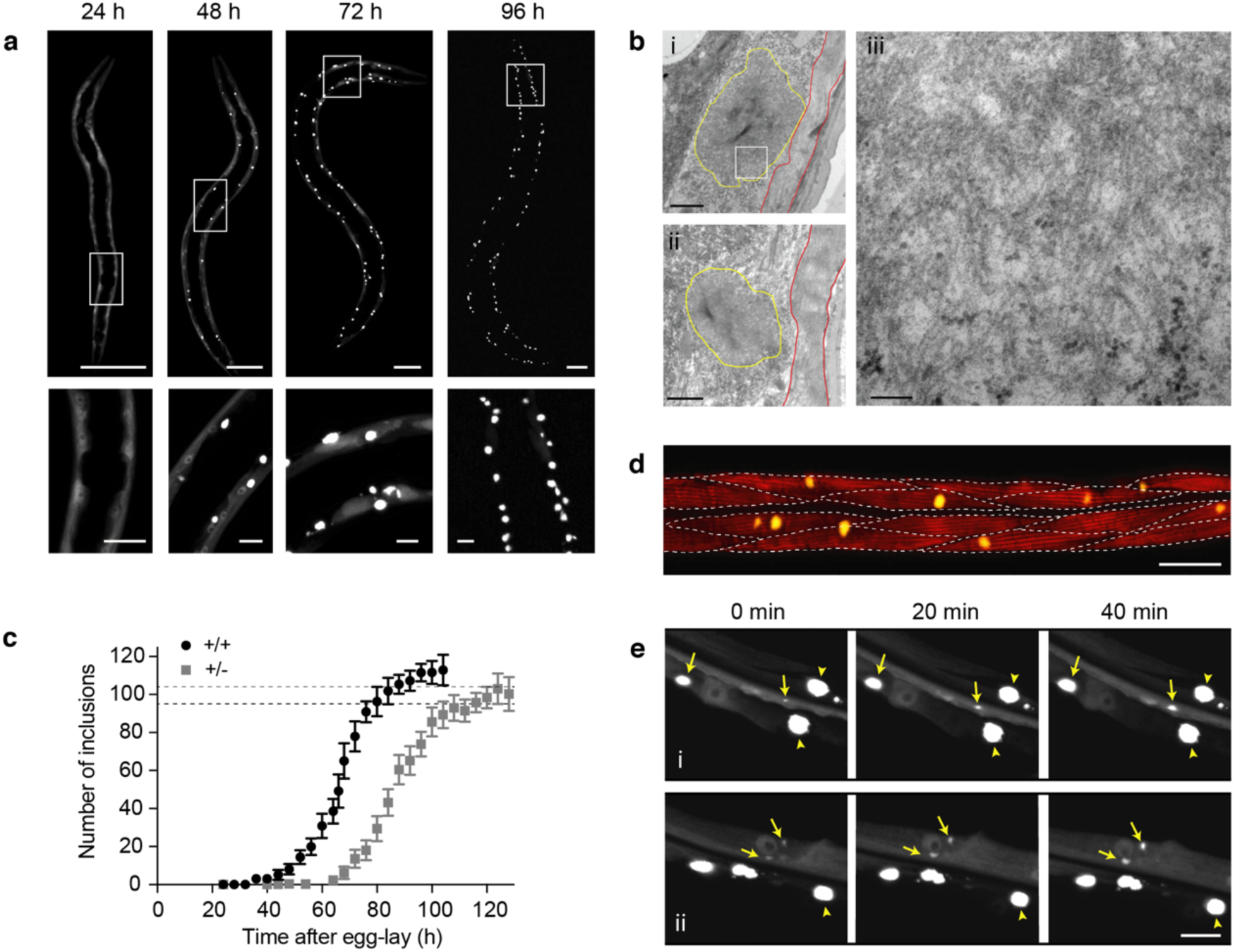
Q40-YFP expressed in *C. elegans* body wall muscle cells displays concentration-dependent, amyloid-like aggregation kinetics with each cell acquiring one inclusion on average. **a**, Confocal images of *C. elegans* strain AM141, progressively displaying bright inclusions of a relatively uniform size and shape in the body wall muscle cells. Lower panels are magnifications of the boxed areas in the upper panels. Scale bars: 50 µm (upper panels) and 10 µm (lower panels). **b**, Transmission electron micrographs of embedded sections of 96h-old animals showing the subcellular localization of the inclusions (yellow outlines) below the muscle sarcomere (red outlines). The right panel shows a higher magnification image corresponding to the boxed region in the upper left panel, displaying a meshwork of fibrils with a typical width on the order of 10 nm. Scale bars: 1 µm (left panels) and 100 nm (right panel). **c**, Average number of inclusions per animal over time for age-synchronized populations of *C. elegans* expressing Q40-YFP in body wall muscle cells for homozygous (+/+) and heterozygous (+/-) animals. The plateau value for the number of inclusions approximates the number of cells in which the protein is expressed, indicated by the dashed lines at 95 for the body wall muscle cells and 104 for the total number of muscle cells in which Q40-YFP is observed. *n* = 12-20 animals for each time point; error bars indicate the standard deviation. **d**, Confocal image of an animal stained with phalloidin around the midpoint of aggregation (66 h) to reveal the muscle filaments. Dashed lines indicate the approximate boundaries between muscle cells, revealing that some have acquired an inclusion by this time, whereas in other cells visible aggregation has not yet taken place. Scale bar: 20 µm. **e**, Panels i and ii: maximum projection confocal images of strain AM141 followed for 40 minutes at around the midpoint of aggregation (64 h). Yellow arrows point to inclusions that are observed to grow during this time, whereas arrow heads indicate mature inclusions that do not change in size. Note that diffuse signal of soluble Q40-YFP is depleted around the mature inclusions. Scale bar: 10 µm.

**Fig. 2:**
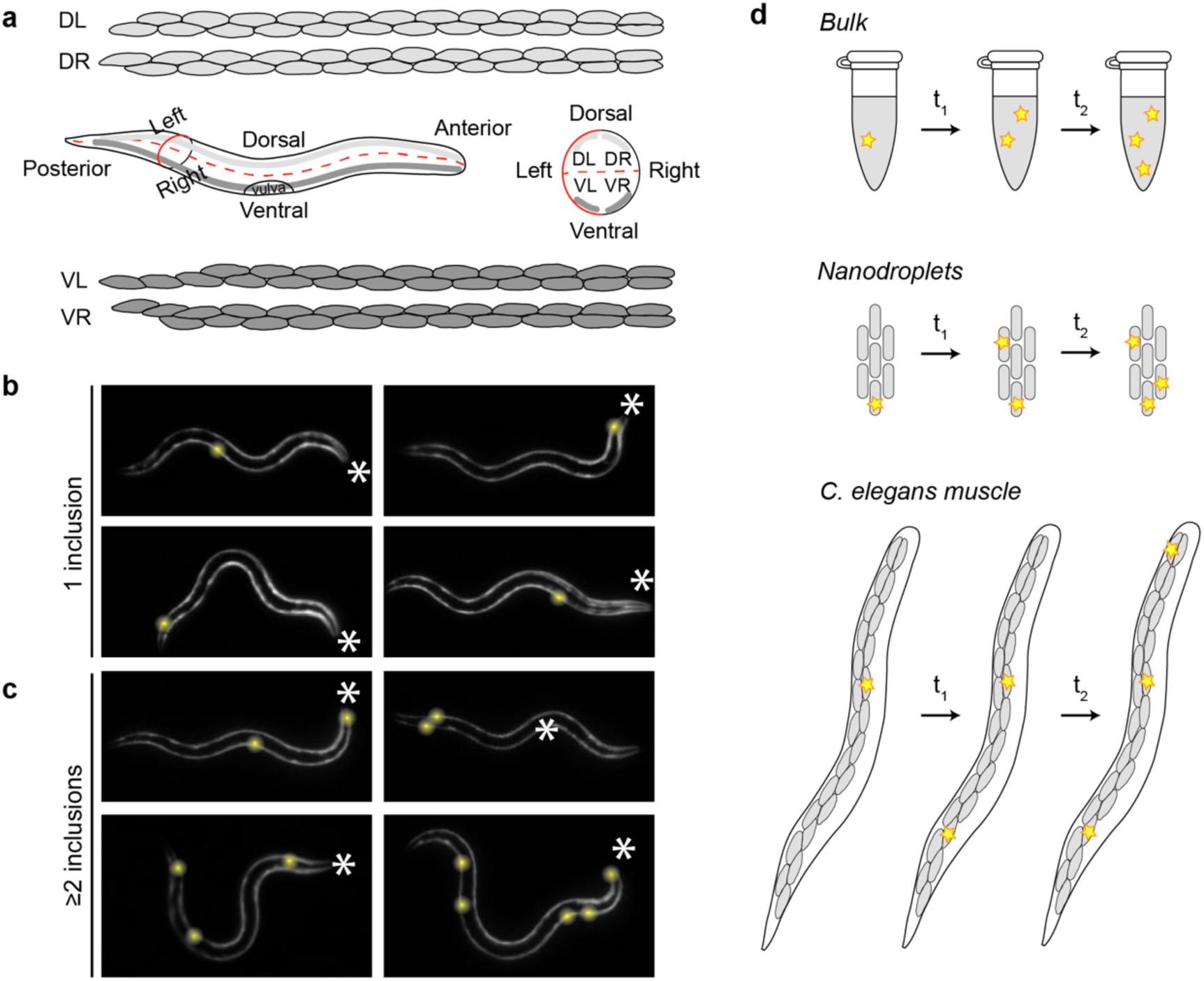
Q40-YFP aggregation occurs stochastically in individual muscle cells. **a**, Illustration showing the distribution of the 95 body wall muscle cells in adult *C. elegans*. Four bundles of muscle cells run along the posterior-anterior axis. The animals typically crawl on their left or right side on solid media, resulting in a superposition of the two dorsal and the two ventral bundles (middle left panel). A schematic view from the anterior side shows the localisation of the four bundles in the different quadrants (middle right panel). Cell shapes were drawn based on wormatlas.org. **b**, 32h-old animals display their first inclusion at any position along the posterior-anterior axis. The asterisks mark the anterior of the animals; inclusions are highlighted by yellow spheres for better visualisation. **c**, 32-h old animals with two or more inclusions support the notion that nucleation is initiated stochastically in individual cells, in the absence of a pattern of spatial propagation from the first to subsequent inclusions. **d**, Cartoon showing the occurrence of aggregation events (yellow stars) in bulk, nanodroplets and in *C. elegans* muscle cells. In a typical test tube reaction (top), all aggregation events take place within the same continuous volume. In nanodroplets (middle) and *C. elegans* muscle tissue (bottom, only one bundle of muscle cells is shown for clarity), the total volume is divided over multiple small volumes, in which the probability of nucleation is low. As a consequence, aggregation will be initiated in different droplets or cells at different points in time. Three aggregation events are shown in each system for clarity, but in reality the number of nucleation events in a given period of time is proportional to the size of the reaction vessel.

Similar to *in vitro* reactions, inclusion formation is concentration-dependent, as evidenced by the comparison of homozygous and heterozygous Q40-YFP animals (Fig. 1c), the latter having 50% of the Q40-YFP gene copies and exhibiting 50% of the fluorescence intensity prior to the onset of aggregation (Extended Data Fig. 3). Despite the different kinetic profiles for the animals expressing the two Q40-YFP protein concentrations, the plateau value for the number of inclusions per animal is very similar (∼112 for homozygous versus ∼100 for heterozygous animals) (Fig. 1c). This number of aggregates corresponds closely to the number of cells in which Q40-YFP is expressed (dashed lines in Fig. 1c). The unc-54 promoter predominantly directs expression to the 95 body wall muscle cells, and in addition strain AM141 displays fluorescence in the eight vulval muscle cells and the anal depressor cell, leading to a total of 104 cells in which Q40-YFP is expressed and aggregation can occur. This result reveals that, on average, each cell acquires one inclusion over the course of the aggregation process. The aggregation curve can therefore be interpreted as the cumulative number of cells in which visible inclusion formation has taken place. Indeed, at the mid-point of the aggregation reaction we find that the majority of cells have either zero or one inclusions, and it is rare to find a cell with two inclusions (Fig. 1d).

**Fig. 3:**
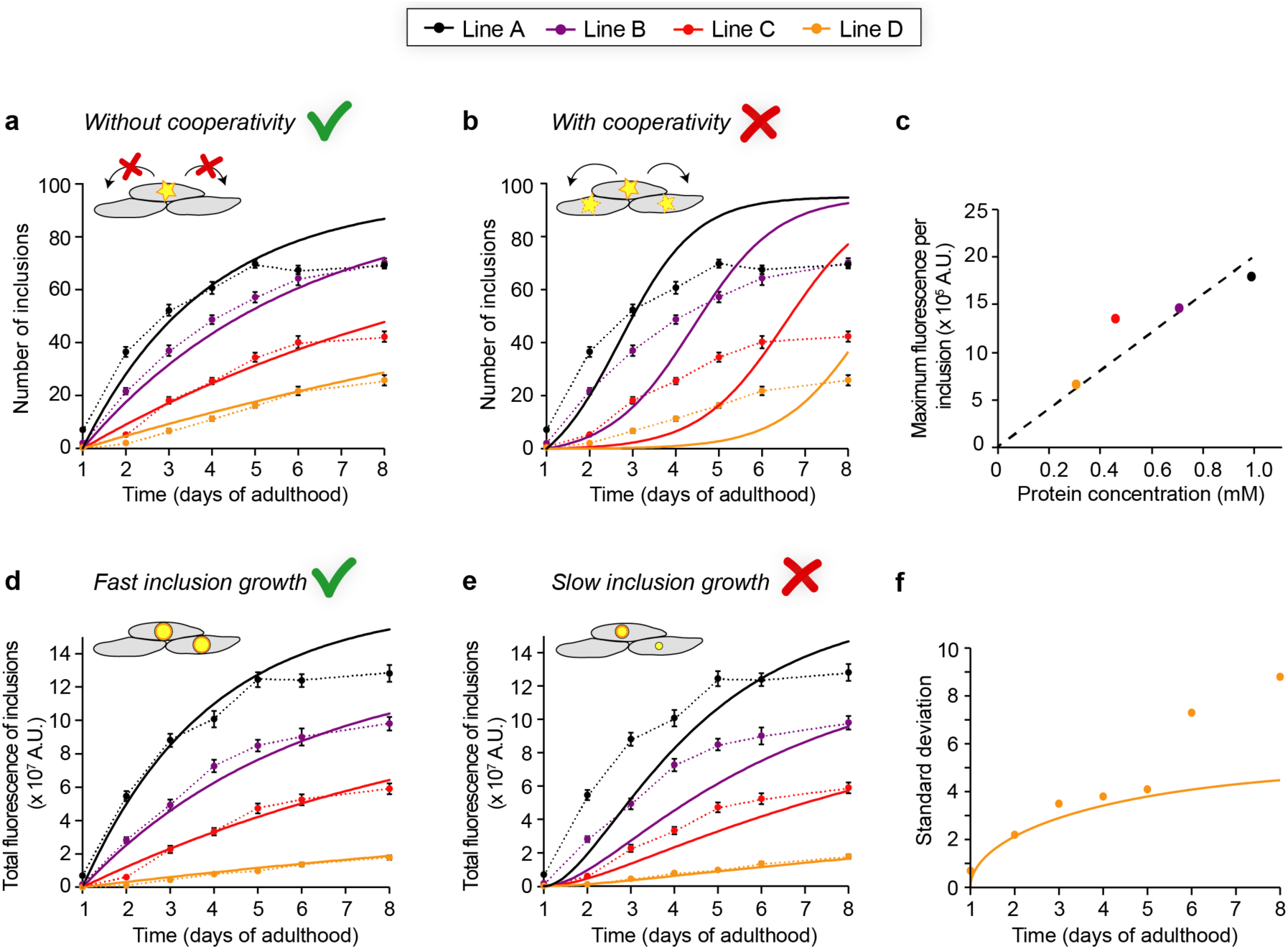
Determining the mechanisms of Q40-YFP aggregation *in vivo* by global fitting of aggregation timecourses at multiple protein concentrations. **a**, Global fit (solid lines) of the average number of inclusions per animal, assuming a constant nucleation rate over time and no cooperativity. The two global free parameters are the nucleation rate and the reaction order. The dashed lines connecting the datapoints in **a**,**b**,**d**,**e** are to guide the eye, and error bars indicate the SEM. *n* = 20 animals per strain and timepoint. **b**, Global fit (solid lines) of the same dataset shown in **a**, but using a model that forces significant cooperativity. Cooperativity is not spatially restricted in this model and can occur between any of the cells. The two global free parameters are the nucleation rate and the reaction order. **c**, The maximum integrated fluorescence intensity per inclusion scales with the intracellular protein concentration for the four Q40-YFP lines. **d**, Prediction of the total aggregate amount based on the fit in **a** and assuming fast inclusion growth on a timescale of hours (solid lines), compared to the observed integrated fluorescence over the inclusions per animal. **e**, Prediction as in **d**, but assuming slow inclusion growth on a timescale of days. **f**, The stochasticity of nucleation accounts for the majority of the experimentally observed standard deviation in line D (*n* = 20 animals). The solid line shows the theoretical prediction based on the nucleation rate for line D, as fitted in **a**. The data shown in this figure are representative of two independent experiments.

The observation that most muscle cells acquire just one polyQ inclusion suggests a rate-limiting nucleation event that initiates the aggregation process. For simple polyQ stretches, nucleation has been proposed to arise from a conformational change of a single molecule^25,26^. However, oligomeric species formed by polyQ-containing proteins have also been observed in several model systems including *C. elegans*^27^, and have been suggested to act as precursors on the pathway towards fibril formation^28–30^. Irrespective of the molecular nature of the nucleation event in our model, this step is required to initiate the formation of visible inclusions through fibril elongation and secondary processes that together amplify the fibril mass. In particular, branching of fibrils or secondary nucleation with slow off-rates of the newly formed nuclei can result in three-dimensional growth of the inclusion. Consistent with this framework, the growth of polyQ inclusions is experimentally observable in the cells of live animals and proceeds at a rapid radial expansion rate of approximately 0.5 µm/h (Fig. 1e). Since mature inclusions typically have a diameter of 3-5 µm, the obtained rate reveals that the inclusions can reach their maximum size on a timescale of several hours. Diffuse signal corresponding to soluble Q40-YFP is no longer observed surrounding mature inclusions (Fig. 1e), indicating that inclusion growth is limited by the available amount of protein in each cell.

### Nucleation occurs stochastically in individual muscle cells

The gradual increase of the number of cells in which an inclusion has formed (Fig. 1c,d) implies that the initiation of protein aggregation occurs at different moments in time in individual cells. One possible explanation for this observation could be biological variation that makes some cells more prone to undergo aggregation than others. However, aside from the vulval muscle cells and the anal depressor muscle, the 95 *C. elegans* body wall muscle cells (Fig. 2a) are highly similar in their lineage relationship, anatomical function and gene expression profiles^31,32^. Consistent with the cells being physiologically identical and having equal aggregation propensities, the first inclusion observed for each of a set of animals appeared randomly among the cells along the posterior-anterior axis (Fig. 2b, Extended Data Fig. 4a,c,d).

A second possible hypothesis for the different aggregation onset across cells is that nucleation is first randomly triggered in one cell, after which aggregation spreads to neighbouring cells. The spreading of aggregation throughout the brain by means of cell-to-cell transfer of aggregate seeds has been proposed to contribute to the pathology of several neurodegenerative diseases, including Huntington’s^33,34^. However, we do not find evidence for a spatial correlation between the first and subsequent inclusions (Fig. 2c, Extended Data Fig. 4b), suggesting that aggregation is triggered by independent nucleation events in the corresponding cells in *C. elegans*. The absence of direct aggregate transfer between adjacent cells is in agreement with a recently published observation that Q40-RFP expressed in *C. elegans* body wall muscle does not spread to neighbouring tissues^35^, and is further verified below using kinetic analysis.

These observations motivate the question whether the inclusions form at different times in each cell because of the intrinsic stochasticity of the nucleation process in small volumes (Fig. 2d). In bulk *in vitro* experiments, nucleation events occur with high frequency throughout the continuous, microlitre-sized reaction volume. In contrast, these events become very rare when one considers individual picolitre-sized volumes, such as that of a cell^13–15^. The intrinsic nucleation properties of the molecules remain the same when a given reaction volume is divided into multiple small volumes, but one cannot predict in which of the small volumes a given nucleation event will occur. In the *C. elegans* muscle tissue, aggregate nuclei will thus appear randomly distributed in a cell-by-cell fashion over time (Fig. 2d). The probability of a second nucleation event occurring within the same cell is low, especially given the rapid speed of inclusion growth which quickly depletes the surrounding soluble protein (Fig. 1e), hence decreasing the probability of another nucleation event. Altogether, our observations are in agreement with a model in which Q40-YFP aggregation in *C. elegans* body wall muscle cells is governed by stochastic nucleation in individual cells, followed by inclusion growth.

### Quantitative analysis of *in vivo* protein aggregation kinetics

To consolidate these findings of stochastic nucleation and inclusion growth in a quantitative manner, we turned towards the application of chemical kinetics. Kinetic analysis has transformed the mechanistic understanding of protein aggregation *in vitro* and allows macroscopic measurements of aggregation processes to be connected with their microscopic mechanisms. Global fitting of *in vitro* data depends on the availability of a range of protein concentrations^4^, which we can achieve *in vivo* through the use of several different strains with different expression levels. Moreover, under *in vitro* conditions, the volume within which the aggregation reaction takes place is typically constant throughout the aggregation timecourse. In strain AM141, inclusion formation begins during larval stages, and we found that the marked increase in muscle cell volume during development (ca. 50-fold from embryo to adult, Extended Data Fig. 5a) dominates the aggregation curve for this strain, given that the probability of nucleation is expected to scale linearly with cell volume (Extended Data Fig. 5). Hence, we generated a new series of four *C. elegans* strains (designated A-D) that express a range of Q40-YFP levels under the control of the unc-54 promoter. The reduced protein concentrations compared to strain AM141 restrict nucleation during developmental stages, and instead the new lines show an onset of inclusion formation at early adulthood, when the muscle cells are close to reaching their maximum volume (Extended Data Fig. 5a).

To monitor the aggregation timecourse for all four strains in parallel, we employed a high-throughput confocal imaging platform with semi-automated image analysis (Fig. 3, Extended Data Fig. 6, Extended Data Fig. 7, see Methods). This method yields very similar numbers compared to those obtained by manual inclusion counting (Extended Data Fig. 6e), but has the advantage of being more quantitative and unbiased. The intracellular protein concentrations were quantified from western blots (Extended Data Fig. 8, see Methods), and as expected strains expressing higher protein concentrations are associated with faster kinetics (Fig. 3, Extended Data Fig. 7a). We monitored the animals from day 1 up to day 8 of adulthood, since few animals remained healthy and alive beyond this timepoint (not shown). Fluorescence intensities were proportional to the protein concentrations as determined by western blot analysis (Extended Data Fig. 8d), and remained approximately constant throughout the timecourse of the experiment (Extended Data Fig. 7b).

In order to define the Q40-YFP aggregation reaction in *C. elegans* muscle cells by means of the underlying nucleation rate constant and the corresponding reaction order, we developed a model of stochastic nucleation with explicit dependence on the protein concentration (see Methods, equation 1). In this model, we assume that the time between nucleation and the appearance of an observable aggregate is short compared to the timescale of the measurement, which is supported by the rapid growth of detectable aggregates (Fig. 1e), and that the aggregation behaviour is deterministic once a nucleus has formed. Furthermore, we chose to fix the plateau value at 95 following our earlier observation that, on average, each body wall muscle cell acquires one inclusion. We first fitted the numbers of inclusions for the four *C. elegans* lines to a model that assumes a constant nucleation rate over time (Fig. 3a), and extracted a reaction order of 1.6 with respect to the protein concentration and a nucleation rate constant of 6 × 10^−13^ M^-0.6^ s^-1^. At an intracellular protein concentration of 1 mM, as in line A, this rate constant corresponds to a nucleation rate of 7 × 10^−18^ M s^-1^, or 0.01 molecules h^-1^ per cell. The significant dependence of the nucleation rate on the protein concentration, the former being proportional to the 1.6^th^ power of the latter, implies that a nucleus size of less than 2 is unlikely. The fits to this model generally represent the experimental data well, with the exception of the last timepoints in the highest expressing line A animals. It is unclear why the number of inclusions ceases to increase beyond day 5 of adulthood for line A, and we speculate that it may be due to positive selection for animals that remain healthy, or due to the increased proteasomal activity that has been observed in ageing *C. elegans*^36^, leading to degradation of the remaining soluble protein.

As noted above, there is currently no evidence to support direct transfer of aggregated species between neighbouring cells in *C. elegans* polyQ models (Fig. 2c, Extended Data Fig. 4b)^35^. Communication through the regulation of proteostasis mechanisms, however, is not spatially restricted and can occur between all cells in a tissue, and even between different tissues within an organism^37^. This phenomenon could lead to cooperativity in the aggregation behaviour, even in the absence of a spatial correlation in the appearance of aggregates. To test if such a phenomenon plays a significant role, we extended our mathematical model to include a cooperative mechanism by which the presence of cells with an inclusion increases the aggregation probability in other cells. The predicted kinetic profiles display initial upwards curvature and do not reproduce the data well (Fig. 3b), leading us to conclude that aggregation in the *C. elegans* body wall muscle cells occurs independently in individual cells with a constant nucleation rate.

The image analysis furthermore allowed us to extract measures for the aggregate mass per inclusion and per animal, based on the integrated fluorescence intensity of the thresholded inclusions. We observed that the average total fluorescence per inclusion converges to a plateau value for each strain (Extended Data Fig. 7c), and that these values are approximately proportional to the protein concentration (Fig. 3c). This finding is in line with our previous notion that, in each cell, the inclusion grows until all diffuse protein is depleted (Fig. 1a,e). Using the average maximum fluorescence for the inclusions in each strain combined with the fits obtained from the number of aggregates, we predicted the development of the total fluorescence intensity of the inclusions per animal under different assumptions about the timescale of inclusion growth. Assuming rapid growth (reaching the maximum inclusion size within approximately 4 hours), the predictions closely match the experimentally obtained values as determined by the integrated fluorescence over the inclusions per animal (Fig. 3d). Predictions for slow growth (inclusions reaching their maximum size in approximately 3 days), on the other hand, are not in agreement with the experimental data (Fig. 3e).

The stochastic nature of nucleation in small volumes not only leads to cell-to-cell differences within a single animal, but may also play a role in animal-to-animal variation given the finite number of cells within each animal. Inter-animal differences in the number of inclusions were previously attributed primarily to biological variation^23^. However, simulations for a 95-cell system, based solely on our fits of the data in Fig. 3a, show that stochasticity causes a considerable standard deviation in the number of inclusions in a population of animals (Fig. 3f, Extended Data Fig. 9). We find that the predictions are in remarkable agreement with the experimental values obtained for the lowest expressing strain D up to day 5 of adulthood (Fig. 3f), and present a lower boundary for the experimental values at later timepoints and in higher expressing strains, for which measurement errors and biological variation presumably make additional contributions (Fig. 3f, Extended Data Fig. 9).

### Nucleated polyQ aggregation mechanism is conserved across experimental systems

Having established a biophysical framework for polyQ aggregation in *C. elegans* muscle cells (Extended Data Fig. 10), we next compare our results to other experimental systems, and examine the applicability to the human brain. The known human polyQ diseases are associated with different, unrelated proteins, carrying different flanking regions adjacent to the polyQ expansion that have been shown to modulate aggregation kinetics as well as toxicity in a variety of experimental systems^38,39^. However, all polyQ diseases have a pathogenic threshold starting at an expansion of around 40 glutamine residues^40^, and this threshold is conserved across experimental systems including *C. elegans*^23^.

The first key finding emerging from our work is that Q40-YFP aggregation in each *C. elegans* muscle cell is governed by stochastic nucleation, followed by growth of the inclusion on a timescale that is fast compared to that of nucleation (Fig. 1e, Fig. 3d,e). Notably, typically one inclusion per cell is observed in cell culture^41–43^, in mouse models of polyQ diseases^22,44,45^ and in patient material^19,21^, consistent with the existence of a rate-limiting nucleation event followed by fast inclusion growth as observed in this study.

The second key finding is that it is possible to quantify the nucleation rate constant and reaction order from global fitting of *in vivo* kinetic data (Fig. 3a), allowing for a comparative analysis across experimental systems. The nucleation step for Q40-YFP aggregation in the *C. elegans* body wall muscle cells shows a significant dependence on the protein concentration, with a reaction order of approximately 1.6. This value is similar to that found in previous work on several polyQ proteins both *in vitro*^5^ and in cell culture^6^, raising the possibility of a conserved nucleation mechanism. The timescale of nucleation as demonstrated in cultured cells is comparable to that in *C. elegans*, but the intracellular protein concentrations in these cells (several µM) were approximately two orders of magnitude lower than for *C. elegans* (several hundred µM)^6^. Therefore, the rate constants of the reaction differ significantly, being orders of magnitude higher in the cells. Our quantitative framework can be applied in future studies to clarify whether this discrepancy is due to the use of different polyQ constructs, the difference between the respective cell types, between isolated cells and a living organism, or a combination of these factors.

Our third key finding is that nucleation occurs independently in each body wall muscle cell, i.e. the extent to which aggregation in one cell increases aggregation in other cells is negligible (Fig. 3a,b). The observation that each cell operates essentially as an individual entity in which nucleation occurs stochastically is consistent with cell culture data^6^, but is remarkable in the context of a multicellular animal. We do not find evidence for spreading of Q40-YFP aggregates in *C. elegans* (Fig. 2c), in contrast to previous observations for a yeast prion protein^46^ and for α-synuclein^35^, suggesting this is not due to a limitation of this animal model. The collapse in proteostasis that is thought to underlie age-associated protein aggregation^36,47–49^, on the other hand, would also lead to cooperative aggregation kinetics. Possibly, the protein concentrations in our *C. elegans* models are associated with nucleation rates that are sufficiently high to dominate over cooperative effects, such as those mediated by proteostasis. The concentration of a polyQ protein such as huntingtin in the human brain, however, is about four orders of magnitude lower^50^. Considering the nucleation rate and reaction order that we find in *C. elegans*, and using a cell volume of 4 pL, we calculate that nucleation would only occur in about 0.5% of cells in the human brain over a period of forty years. However, a fraction of ca. 20-30% of neurons has been reported to contain inclusions in patient material^21^, suggesting that aggregation kinetics in the human brain are faster than we extrapolate from our results obtained in *C. elegans*. Moreover, no inclusions were found in a presymptomatic individual that carried the disease allele^21^, suggesting that protein aggregation is only initiated in mid-life when symptoms appear. A significantly cooperative model may therefore be more appropriate to describe the progression of human polyQ aggregation around the pathogenic threshold.

In conclusion, our results show that the same biophysical principles that govern protein aggregation in small volumes *in vitro*, namely spatial confinement and stochastic nucleation, remain the dominant driving forces in an *in vivo* system. We anticipate that our modelling approach will be applicable to analyse the features of amyloid-like protein aggregation in wide range of biological systems, including human tissues affected by protein misfolding diseases.

## Acknowledgements

We thank the Biological Imaging Facility, High-Throughput Analysis Lab, BioCryo Facility and Keck Biophysics Facility and the Lackner lab at Northwestern University for instrument use and technical assistance. We are grateful to Renée Brielmann for *C. elegans* micro-injections and to Charlene Wilke for freeze substitution and assistance with TEM. This work was supported by grants from the NIH (National Institute on Aging R56AG059579, R37AG026647, RF1AG057296 and P01AG054407) and the Daniel F. and Ada L. Rice Foundation to R.I.M..

## Author contributions

T.S., G.M., T.C.T.M., M.V., T.P.J.K. and R.I.M. conceived the study and interpreted data. T.S. and R.I.M. designed experiments, and T.S. performed experiments and analysed data. G.M. and T.C.T.M. developed the mathematical models and performed fitting with contributions from M.V. and T.P.J.K.. T.S. drafted the manuscript with input from R.I.M. and G.M., and all authors edited the manuscript.

## Competing interests

The authors declare no competing interests.

## Correspondence and requests for materials

should be addressed to T.S. or R.I.M.

## Methods

### *C. elegans* methods and strains

Nematodes were maintained on nematode growth media (NGM) seeded with *Escherichia coli* OP50 at 20°C using standard methods^51^. Age-synchronized populations of *C. elegans* were generated by allowing adult animals to lay eggs on NGM plates for a period of 1-2 h. For long-term experiments, animals were transferred daily to fresh NGM plates to separate the adults from their offspring.

Strains used in this study are:

N2

AM134, rmIs126 [unc-54p::Q0::YFP] X

AM141, rmIs133 [unc-54p::Q40::YFP] X

AM1228, rmIs404 [unc-54p::Q40::YFP] *“line A”*

AM1229, rmIs404 [unc-54p::Q40::YFP] *“line B”*

AM1230, rmIs404 [unc-54p::Q40::YFP] *“line C”*

AM1231, rmIs404 [unc-54p::Q40::YFP] “*line D”*

### Generation of transgenic *C. elegans* strains

Strains AM1228, AM1229, AM1230 and AM1231 were generated by micro-injection of plasmid pPD30.38 unc-54p::Q40::YFP, the same plasmid that was used to create strain AM141^23^. Extrachromosomal arrays were integrated by UV irradiation, and the strains were backcrossed five times with wild-type N2 worms to remove background mutations.

### Fluorescence imaging of inclusions

Manual counting of inclusions was performed using a fluorescence stereomicroscope (Leica MZ16FA) at 115x magnification, while the animals freely crawled on seeded NGM plates. For each time point, 12-20 animals were counted from the same age-synchronised pool. Wide-field fluorescence images of homozygous and heterozygous animals were acquired using a Leica M205FA stereomicroscope equipped with a Hamamatsu C10600 digital camera at 100x or 160x magnification.

For semi-automated inclusion counting, 20 animals for each strain and timepoint were picked into one well of a 384-well plate filled with M9 buffer (5.8 g/L Na_2_HPO_4_, 3 g/L KH_2_PO_4_, 0.5 g/L NaCl, 1 g/L NH_4_Cl) supplemented with 25 mM NaN_3_ as an anaesthetic. The animals used for the different timepoints in one experiment were derived from the same synchronized egg-lay, and two independent timecourse experiments were performed. The timepoint for day 1 of adulthood was taken 72h after egg-lay, when the animals had typically started to lay eggs, and subsequent timepoints were taken at ∼24h intervals. Confocal imaging was performed on the ImageXpress high-content imaging system (Molecular Devices) equipped with a 20x objective. 25 maximum projection images were recorded per well and automatically stitched using the instrument software. Artefacts resulting from residual movement of the animals were manually corrected. The resulting images were thresholded in ImageJ after which the inclusions were selected using the ‘analyse particles’ function with a minimum size of 1 µm^2^, allowing the numbers of inclusions and the total fluorescence intensity of the inclusions in each worm to be determined.

### Phalloidin staining

Phalloidin staining was performed by fixing animals in 4% para-formaldehyde in phosphate buffered saline (PBS) for 30 minutes at room temperature, followed by washing them in PBS + 0.05% Tween-20, and treatment with 5% beta-mercaptoethanol in 134 mM Tris-HCl pH 6.8 + 1% Triton-X100 at 37°C and 100 rpm overnight. After washing in PBS + 0.05% Tween-20, fixed animals were incubated in 1% Alexa 633-conjugated phalloidin in PBS + 0.5% Triton X-100 + 1% BSA. After washing in the same buffer without phalloidin, animals were mounted on glass slides with Fluoromount-G mounting media (Thermo Fisher Scientific) and imaged on a Leica SP8 confocal microscope equipped with 40x oil objective.

### Confocal imaging of inclusion growth

Confocal imaging was performed on a Leica SP8 microscope. To determine inclusion growth rates, live animals were immobilized on 3% agarose pads in a drop of M9 with 2 mM levamisole as a mild anaesthetic, and imaged within a period of 1 hour using a 40x oil objective. The rate of inclusion formation under these conditions was confirmed to be similar to that in worms freely crawling on NGM plates, with on average 5-6 new inclusions formed per hour in strain AM141 around the mid-point of aggregation.

### Quantification of cell volume and protein concentrations

The total volume of the muscle cells in adult animals was determined by confocal imaging on a Leica SP8 using a 10x air objective. Cell volumes of developing animals were determined using the 40x or 20x oil objectives. The images were deconvolved using Huygens software, and ImageJ was used to threshold the images and determine the total fluorescent volume in each animal using the Voxel Counter plugin. The average of ten animals was taken for each condition, and the volume per cell obtained by dividing this number by 104, to account for the number of cells in which the protein is expressed under the unc-54 promoter (95 body wall muscle cells, 8 vulval muscle cells and anal depressor cell).

To determine the protein content per animal, a total of 30 animals at the first day of adulthood were picked into M9 buffer for each strain, snap frozen in liquid nitrogen and stored at -80°C until further processing. After addition of 5x SDS Sample Buffer (62.5 mM Tris-HCl pH 6.8, 20% glycerol, 2% SDS, 5% beta-mercaptoethanol, 0.1% bromophenol blue), samples were boiled, and the total volumes were loaded onto SDS PAGE gels. Recombinant EYFP (Ray Biotech) was used as a standard to quantify the amounts of protein in each lane, and the Western blots were probed with JL-8 anti-GFP antibody (Takara Bio) and scanned on an Azure c600 (Azure Biosystems). Three biological replicates were averaged to obtain the final protein amount per worm. This value was converted to intracellular protein concentration using 27 kDa molecular weight for YFP, and 99 pL for the total volume of the cells in which the protein is expressed at the first day of adulthood.

Relative protein amounts in developing AM141 animals were determined from fluorescence images acquired on the ImageXpress confocal platform (Molecular Devices). Relative intracellular protein concentrations were obtained by dividing the integrated fluorescence intensity per animal by the total fluorescent volume per animal determined as described above.

### Transmission electron microscopy (TEM)

96h-old AM141 animals were subjected to high-pressure freezing with a Leica HPM100 using *E. coli* OP50 for freeze protection. Samples stored in liquid N_2_ were transferred to a precooled Leica EM-AFS2 unit with freeze substitution processor (FSP) for automated solution exchanges. The freeze substitution solution was 0.01% OsO4, 0.1% KMnO4 in 95% anhydride acetone and substitution began at -90°C for 24 hours. The temperature was increased at a rate of 6°C/h to -50°C. Samples remained at -50°C for 6 hours. Four 10 minute washes using 95% acetone took place at -50°C. The second freeze substitution solution of 0.1% uranyl acetate in 95% acetone ran for 3 hours with a temperature increase to -40°C over 3 hours, followed by four 10 minute acetone washes. The temperature increased to -30°C for 2 hours (5°C/h) before three 10 minute washes using 95% ETOH. For resin embedding, Lowicryl HM20 was infiltrated at -30°C with the following schedule: 70% for 4 hours, 100% for 12 hours, 100% for 4 hours three times, and then polymerized for 48 hours at -30°C using UV light included in the FSP system.

The 80 nm sections were obtained using a Leica Ultracut S and a DiATOME diamond knife. Sections were post-stained with 2% uranyl acetate and lead citrate. TEM images were acquired with a JEOL 1230 GEM TEM at 80 kV, equipped with a Gatan Orius SC 1000 CCD camera.

### Modelling

#### Independent nucleation

The minimal model of stochastic nucleation is derived as follows: The probability of nucleation occurring in any cell is assumed to be constant throughout time and proportional to the monomer concentration to some power, *n*_*c*_, referred to as the reaction order (of nucleation) in the following. We only consider the first nucleation event in a cell, and a cell is either aggregated or not aggregated. The change in the fraction of aggregated cells, *f(t)*, is then given by

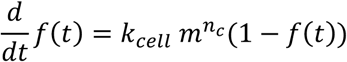

where *k*_*cell*_ is the nucleation rate constant per cell. Solving this and including a factor to convert the fraction of aggregated cells to the average number of aggregated cells, *F(t)= N*_*cells*_ *f*(*t*), gives

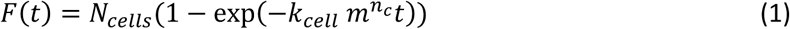

where *N*_*cells*_ is the number of total cells in which aggregates can form, in our case 95. 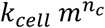 is the total rate (units of inverse time) of nucleation.

#### Cooperativity

To account for the possible presence of cooperativity, we included an additional term in the above model. We assume that the rate at which new aggregates are formed via this cooperative step is proportional to both the fraction of cells already aggregated, as well as the fraction of cells not yet aggregated, yielding the following minimal model of independent and cooperative nucleation

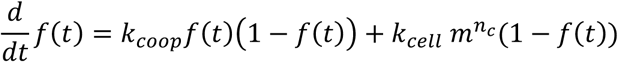

where *k*_*coop*_ is the cooperativity rate. This can be solved for the initial condition of no aggregates to yield

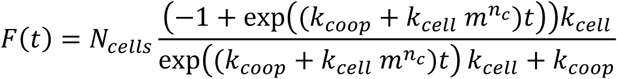

There is no explicit spatial dependence in this model, so it can describe e.g. neighbour-to-neighbour transfer of aggregation transfer, as well as systemic changes due to aggregates that trigger nucleation anywhere in the worm. Note that equation (1) is recovered when *k*_*coop*_ = 0.

The misfits shown in Fig. 3b use a cooperativity rate *k*_*coop*_ = 1 days^-1^. If cooperative nucleation was the only mechanism responsible for aggregate formation, this value for *k*_*coop*_ would lead to a doubling of the number of cells that have an aggregate approximately every 17 hours on average.

#### Including growth

In order to analyse the total fluorescence of the inclusions per animal, we extended the model in equation (1) to include a growth term that describes how the aggregate mass evolves after nucleation. We assume that the aggregate size approaches its maximum value in an exponential manner, which is consistent for example with a linear growth mechanism under constant monomer conditions.

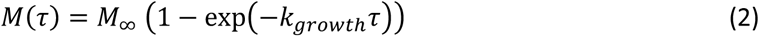

where *M* is the mass of aggregated material in a single cell, *k*_*growth*_ is the growth rate, *τ* is the time since nucleation and *M*_*∞*_ is the mass at the end of the growth reaction. However, as the growth does not affect any of the measurements of the kinetics we recorded here, the specifics of this process are unimportant, and the one parameter of interest is simply the timescale of this process. Equation (2) describes the how the mass evolves given a nucleation event occurred at time *τ* = 0. In order to derive an expression for the total mass of aggregates expected in one animal, this time evolution of the mass after nucleation, equation (2), has to be convoluted with the time evolution of number of aggregates, equation (1), to give

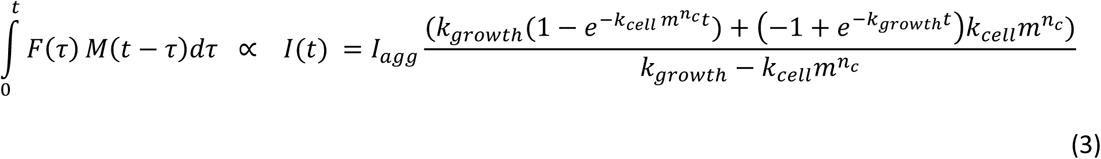

where we have used the assumption that mass is proportional to the total fluorescence of the inclusions, and thus *I(t)* is the total fluorescence of the inclusions, and *I*_*agg*_ is the fluorescence of a single inclusion at its maximal size. In the fits in Fig. 3d we find that the data are well matched as long as *k*_*growth*_ > 20 days^-1^ which is equivalent to the growth reaction being 95% complete in approximately 4 hours. The misfits were produced by setting *k*_*growth*_ = 1 days^-1^ which is equivalent to the growth reaction being 95% complete in approximately 3 days.

#### Effect of volume

A volume that varies over time can be included in our minimal model of nucleation, equation (1), as follows. The volumes are measured at several timepoints, and by inspection the time evolution of the volume is found to follow a sigmoidal shape. Therefore, the functional form

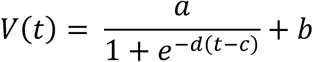

is used to describe the volume. The nucleation rate is assumed to be proportional to the volume of the cell, yielding an updated differential equation for the fraction of cells that have an aggregate as

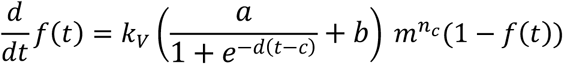

where *k*_*V*_ now denotes a nucleation rate per volume rather than per cell. This can be solved to give

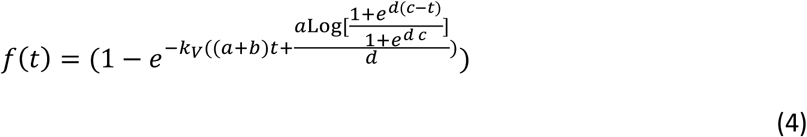

#### Data fitting

Data fitting was performed using a least squares algorithm implemented in our fitting platform, Amylofit^4^. In strains A-D, for the data fitted by equations (1) and (3), the time t=0 was chosen to be day 1 of adulthood as this corresponds to cell volumes and protein expression levels reaching constant values, and virtually no aggregate formation is seen before this timepoint. In strain AM141, fitted by equation (4), the time t=0 was chosen to be 24 h after egg laying as this time corresponds to the protein expression levels reaching a constant value, and the change in volume is taken into account explicitly.

### Statistical analysis

No statistical methods were used to predetermine sample sizes or to analyse datasets.

### Data availability

The data generated during this study are available from the corresponding authors upon reasonable request.

## Notes

### Competing Interest Statement

The authors have declared no competing interest.

