## Extended Data for "Kinetic analysis reveals the rates and mechanisms of protein aggregation in a multicellular organism"

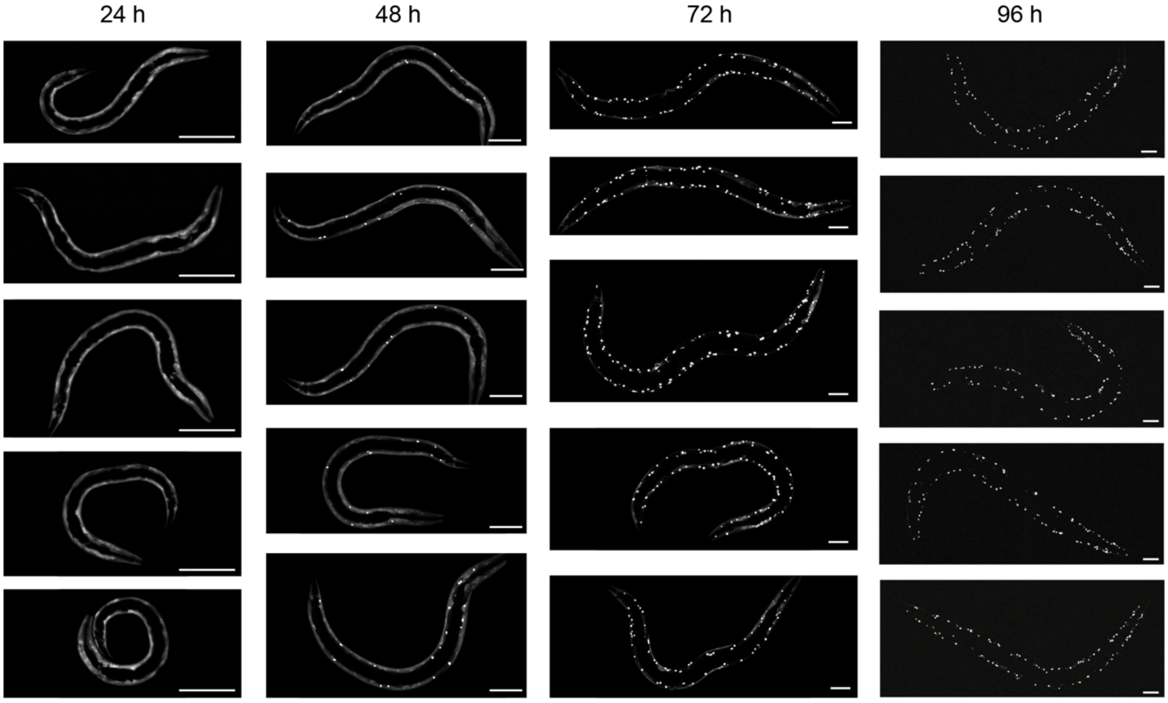


**Extended Data Figure 1: Confocal images of *C. elegans* strain AM141 expressing Q40-YFP in the body wall muscle cells.** Different animals from a time-synchronised population are shown at 24, 48, 72 and 96 h after egg-lay. Scale bars: 50 µm.

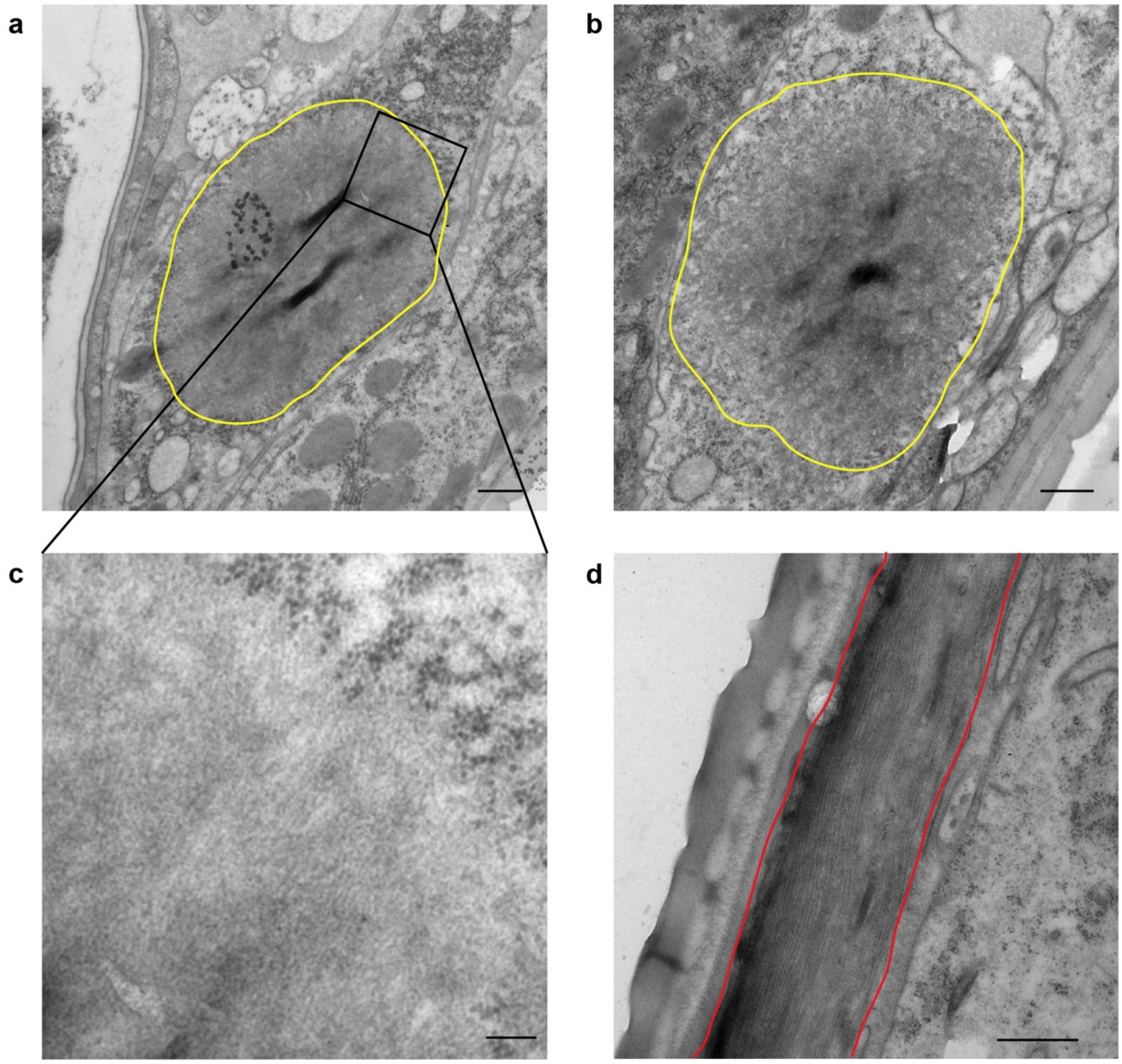


**Extended Data Figure 2: TEM images of *C. elegans* expressing Q40-YFP in the body wall muscle cells.** **a**,**b**, Embedded sections of 96h-old animals showing inclusions outlined in yellow. The muscle sarcomere cannot be seen due to the orientation of these cross-sections. Scale bars: 500 nm **c**, Higher magnification image of the boxed region in panel **a**, allowing fibrillar material to be discerned. Scale bar: 100 nm. **d**, Region of a muscle cell devoid of an inclusion, in which the filamentous structure of the muscle sarcomere can be seen (red outline). Scale bar: 500 nm.

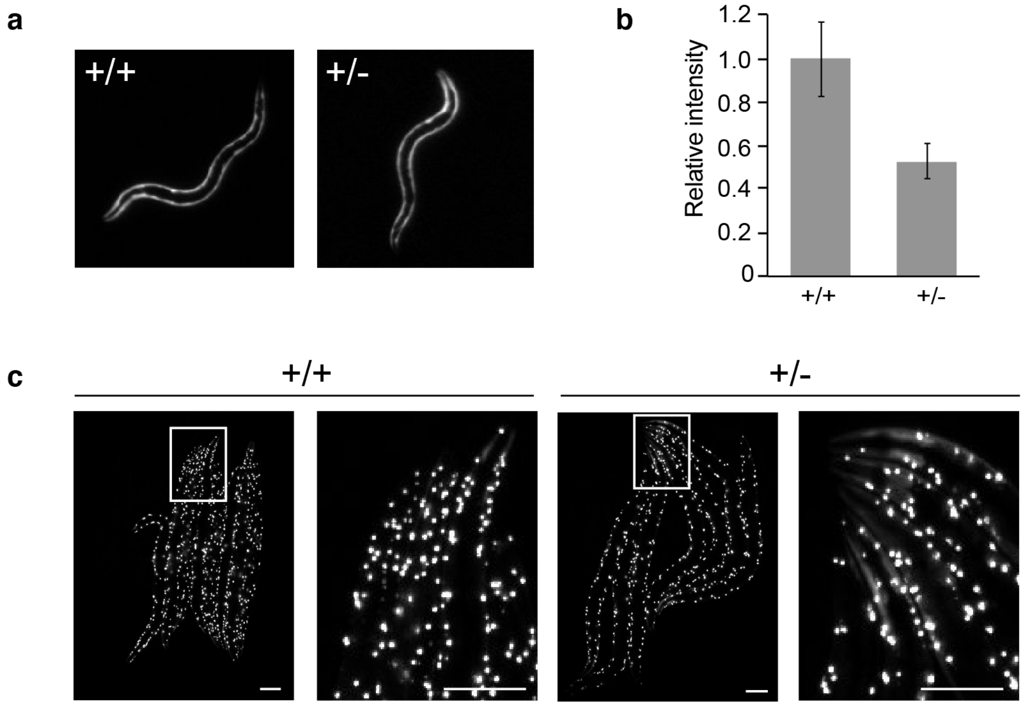


**Extended Data Figure 3: Heterozygous worms contain 50% of the protein amount compared to homozygous worms, yet have the same final distribution of inclusions. a**, Representative fluorescence images of a homozygous (+/+) and a heterozygous (+/-) animal at 27 h past egg-lay, just prior to the onset of aggregation for the homozygous strain. **b**, Quantification of the average fluorescence intensity for ten animals each, relative to homozygous. **c**, Fluorescence images of homozygous (+/+) and heterozygous (+/-) animals taken at the plateau of the aggregation curves (98 h and 128 h, respectively). Diffuse signal has entirely disappeared with the exception of the head region in the heterozygous animals. Boxed regions are magnified in the adjacent panels. Scale bars: 10 µm.

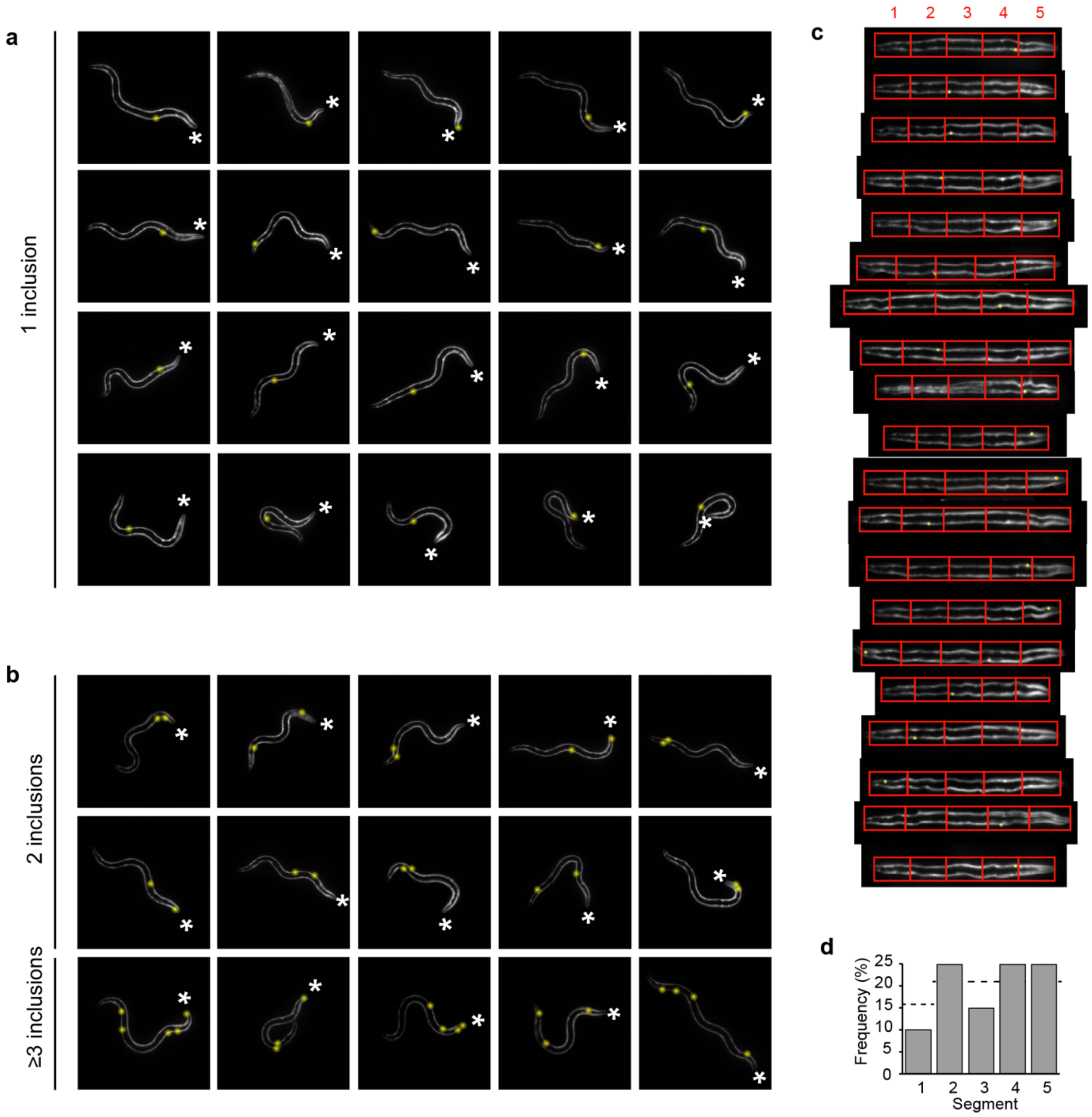


**Extended Data Figure 4: Q40-YFP inclusion formation occurs independently in individual cells. a**, 32h-old animals displaying one inclusion do not show a preferred localisation. **b**, 32h-old animals displaying two or more inclusions do not provide evidence for a spatial correlation between the first and subsequent inclusions. The asterisks mark the anterior of the animals; inclusions are highlighted by yellow spheres. **c**, 20 worm images were straightened in ImageJ and each sectioned into five equal parts along the posterior-anterior axis to determine the location of the inclusion. **d**, The frequency distribution of the location of the first inclusion shows there is no preferred site where aggregation is initiated. The dotted line shows the expected value for an equal distribution with segment 1 containing 15 cells and the other segments 20 cells each.

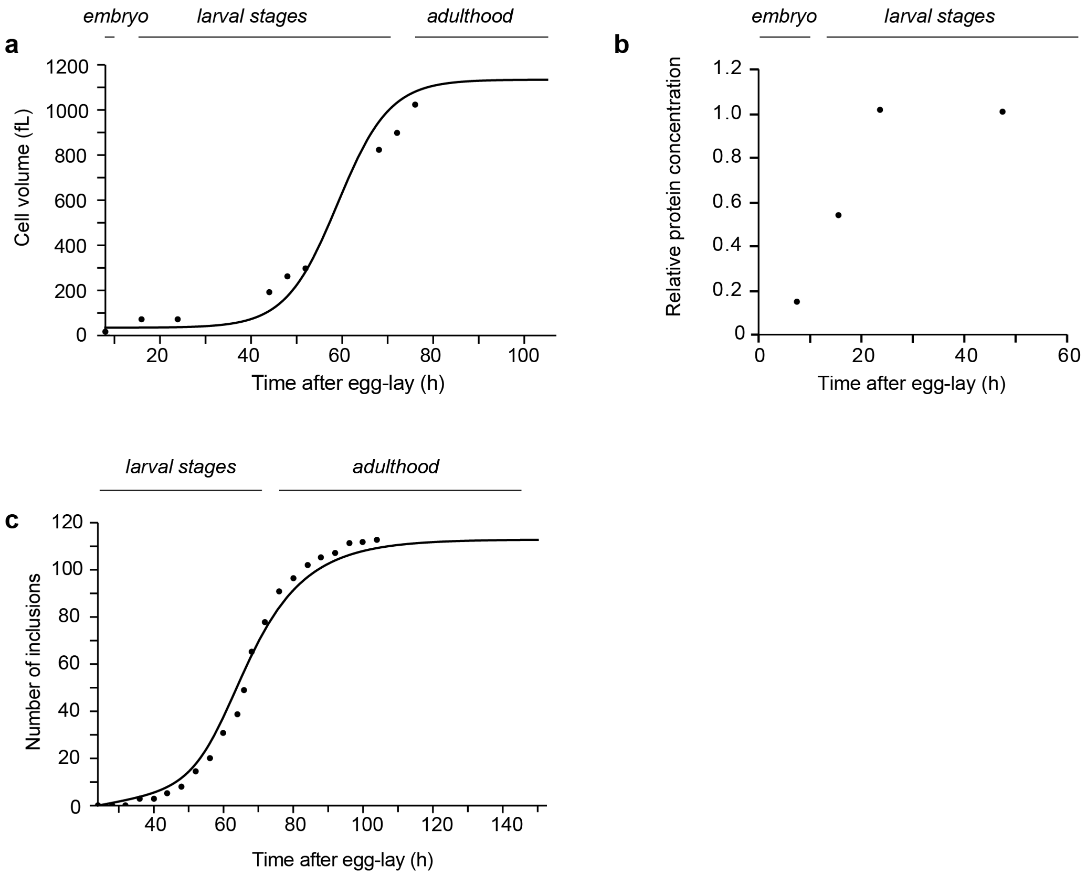


**Extended Data Figure 5: Aggregation kinetics in strain AM141 are largely determined by the increase in cell volume during development.** **a**, Average volume of the body wall muscle cells experimentally determined by confocal microscopy. For timepoints where Q40-YFP localises predominantly to inclusions (beyond 52 h), AM134 animals expressing Q0-YFP were used. AM134 reaches adulthood at around 72 h, compared to 76 h for AM141. Solid line shows a sigmoid used to interpolate the data for use in **c**. *n* = 5-10 animals per timepoint. **b**, Protein expression driven by unc-54 is initiated in late embryonal stages (t = 8 h) and reaches a plateau value around 24 h post egg-lay. Relative protein concentrations were obtained by dividing the fluorescence intensity by the total muscle cell volume at the corresponding timepoint (shown in **a**), and normalising to the highest value. *n* = 20 animals per timepoint. **c**, Simulated aggregation curve (solid line) of strain AM141 (see Fig. 1c, *n* = 12-20 animals per timepoint) assuming that the nucleation rate scales linearly with the cell volume, the time dependence of which is determined in **a**.

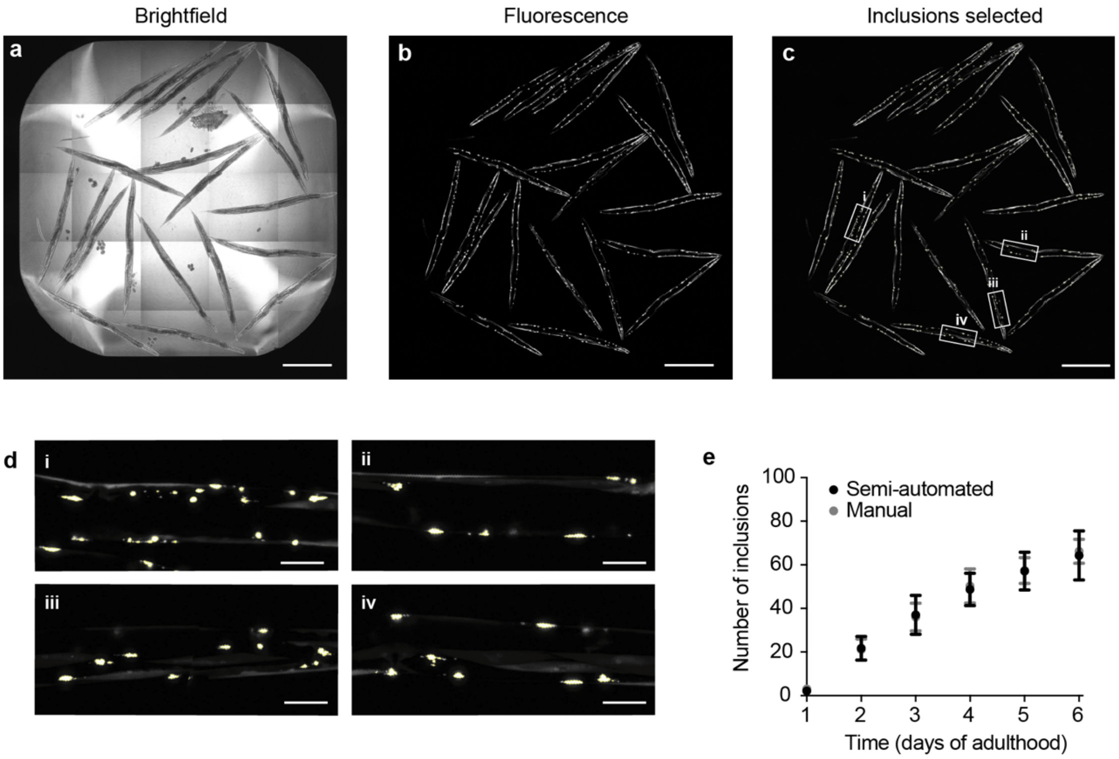


**Extended Data Figure 6: Semi-automated inclusion analysis.** Shown are animals of strain B at day 3 of adulthood. **a**, Brightfield image of one well from a 384-well plate containing 20 animals. The image was recorded as 25 panels using a 20x objective on the high-throughput confocal platform, which were computationally stitched together to obtain the whole well image. **b**, Fluorescence image of the same well. **c**, Overlay of the inclusions selected in ImageJ (yellow outlines) with a minimum area of 1 $\mu m^2$ onto the original fluorescence image. Scale bars in **a-c**: 500 µm. **d**, Magnification of the boxed areas in panel **c** showing the inclusions outlined in yellow. Scale bars: 50 µm. **e**, Comparison of manual versus semi-automated inclusion counting for line B. *n* = 14-20 per timepoint, error bars indicate standard deviation.

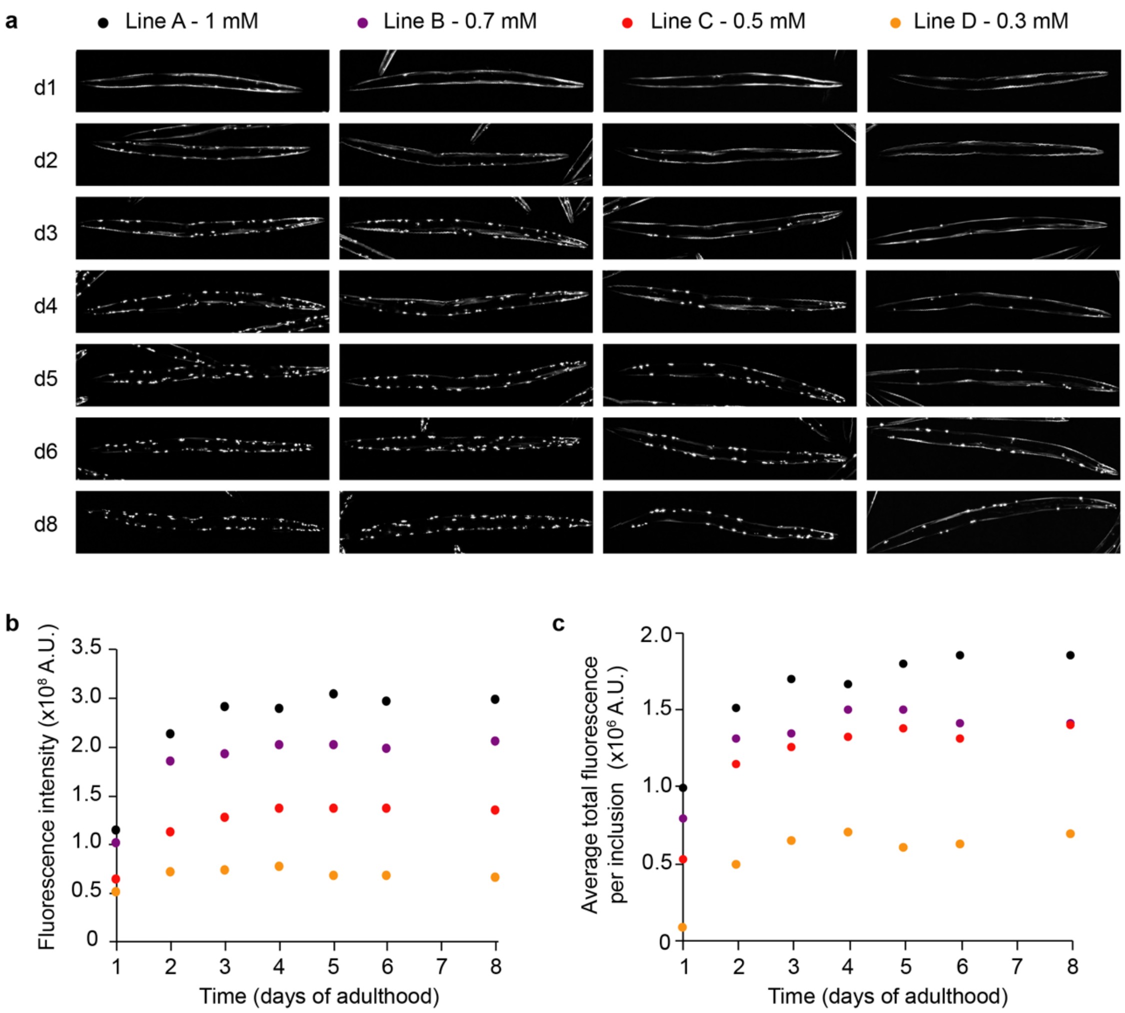


**Extended Data Figure 7: High-throughput imaging of animals of lines A-D showing the development of Q40-YFP inclusions over time. a**, Images were acquired on a high-throughput confocal platform in 384-well plates with 20 animals in each well, from which individual animals are displayed here. The differences in intracellular protein concentrations (from high to low in strain A-D, see Extended Data Fig. 8) are reflected by the aggregation kinetics. **b**, Protein levels are approximately constant throughout adulthood, as seen from the total fluorescence intensity per animal. Only between day 1 and day 2 of adulthood an increase in intensity of approximately 1.7-fold is observed that is associated with a 1.2-fold increase in cell volume (not shown), and hence an estimated 1.4-fold increase in protein concentration. **c**, The average total fluorescence per inclusion rapidly reaches a plateau value for each of the strains, consistent with fast inclusion growth. Note that at day 1 of adulthood, few inclusions are present (Fig. 3a,b), and the average total fluorescence per inclusion is approximately constant from day 2 onwards.

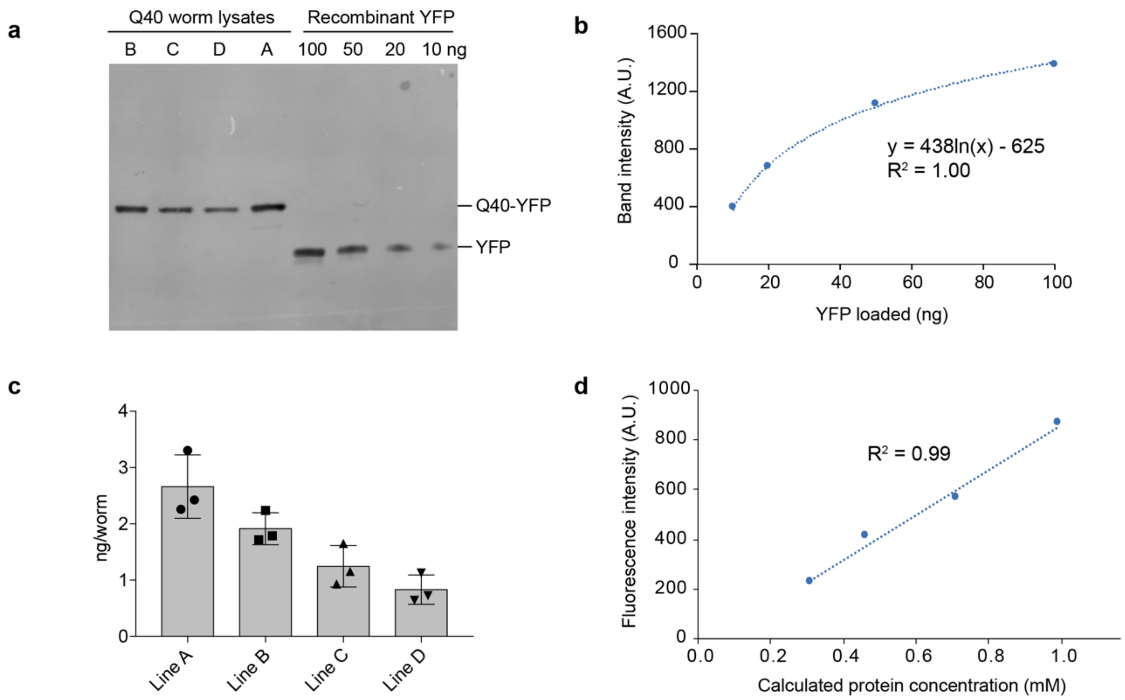


**Extended Data Figure 8: Determination of the intracellular protein concentrations of Q40-YFP. a**, Representative western blot with lysates derived from 30 animals for each strain at day 1 of adulthood, and known amounts of recombinant YFP, probed for YFP. **b**, Quantification of the recombinant YFP band intensities from the blot shown in **a**. The correlation between protein amount and band intensity could be best fitted with a logarithmic function, which was used to calculate the amounts of protein present in the worm lysates. Note that the band intensities of the lysate samples fall in between the YFP calibration points. **c**, Protein amounts in ng/worm for each strain as derived from three independent batches. **d**, The fluorescence intensities of the four strains show a linear correlation with the calculated protein concentrations using the western blot approach.

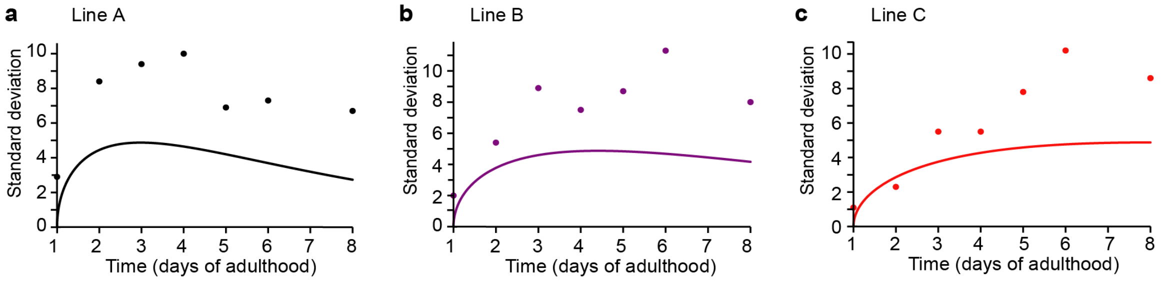


**Extended Data Figure 9: The stochasticity of nucleation sets a lower boundary for the standard deviation of the inclusion numbers in a population of animals. a**, **b**, **c,** Prediction (solid line) and experimental datapoints of the standard deviations for lines A (**a**), B (**b**) and C (**c**). *n* = 20 animals for each strain and timepoint. The predictions are based on the nucleation rate for each strain as determined from global fitting (Fig. 3a).

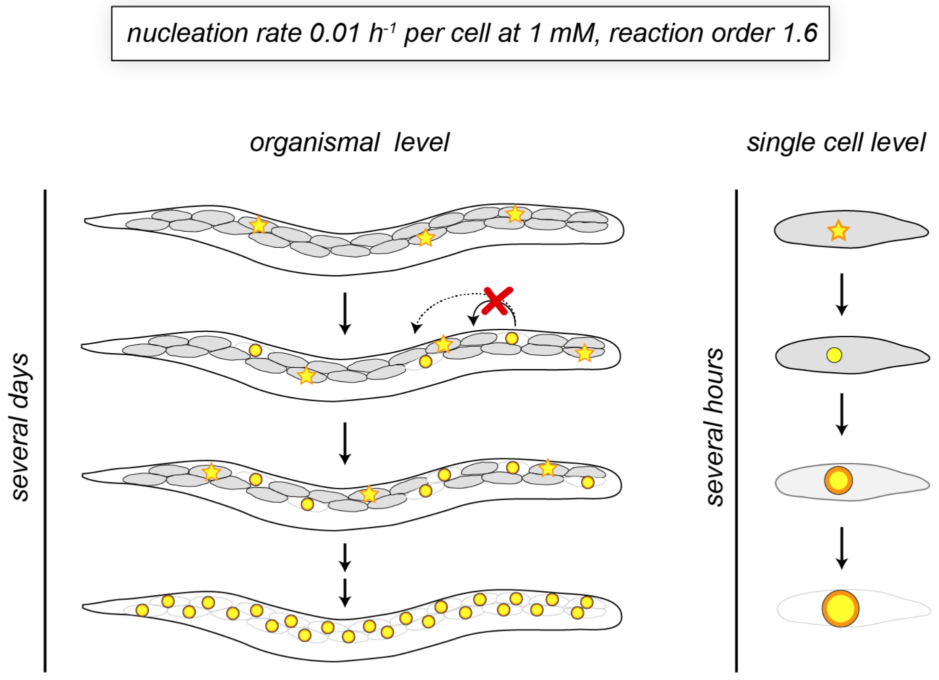


**Extended Data Figure 10: Model of stochastic nucleation and aggregate growth as established in this study.** Q40-YFP aggregation in body wall muscle cells is initiated by stochastic nucleation events that occur independently in each cell at a constant rate of 0.01 $h^{-1}$ per cell at an intracellular protein concentration of 1 mM, with a reaction order of 1.6. Once nucleation has occurred in a particular cell, aggregate growth proceeds rapidly on a timescale of hours until the soluble protein in that cell is depleted.
